# Targeted DNA Sequencing of Cutaneous Melanoma Identifies Prognostic and Predictive Alterations

**DOI:** 10.1101/2024.01.18.576113

**Authors:** Alexandra M. Haugh, Robert C. Osorio, Rony A. Francois, Michael E. Tawil, Katy K. Tsai, Michael Tetzlaff, Adil Daud, Harish N. Vasudevan

## Abstract

**Importance:** Cutaneous melanoma (CM) can be molecularly classified into four groups: *BRAF* mutant, *NRAS* mutant, *NF1* loss, and triple wild type (TWT) tumors lacking any of these three alterations. In the era of immune checkpoint inhibition (ICI) and targeted molecular therapy, the clinical significance of these groups remains unclear. Here, we integrate targeted DNA sequencing with comprehensive clinical follow-up in CM patients.

**Objective:** To explore how molecular features and tumor mutational burden (TMB) impact outcomes in patients with cutaneous melanoma.

**Design:** This was a retrospective cohort study that assessed clinical and molecular features from patients with localized or metastatic CM who underwent targeted next-generation sequencing as part of routine clinical care.

**Setting:** Tertiary referral, comprehensive NCI cancer center from 2013 - 2023.

**Participants:** A total of 254 patients with CM who had a CLIA certified targeted sequencing assay performed on their tumor tissue were included

**Exposure:** A CLIA certified targeted sequencing assay was performed as standard of care on 254 patients with CM treated at a single institution.

**Main Outcome:** *NRAS* mutation correlated with significantly worse overall survival compared to other TCGA driver groups. Elevated TMB correlated with improved progression-free survival on combination checkpoint inhibition (anti-PD1 plus anti-CTLA4).

**Results:** Of 254 patients with cutaneous melanoma, 77 were *BRAF* mutant (30.3%), 77 were *NRAS* mutant (30.3%), 47 were *NF1* mutant (18.5%), 33 were TWT (13.0%) and the remaining 20 (7.9%) carried mutations in multiple driver genes (*BRAF/NRAS/NF1* co-mutated). The majority of this co-mutation group carried mutations in *NF1* (n=19 or 90%) with co-occurring mutations in *BRAF* or *NRAS,* often with a weaker oncogenic variant. Consistently, *NF1* mutant tumors harbored numerous significantly co-altered genes compared to *BRAF* or *NRAS* mutant tumors. The majority of TWT tumors (n=29, 87.9%) harbored a pathogenic mutation within a known Ras/MAPK signaling pathway component. Of the 154 cases with available TMB data, the median TMB was 20 (range 0.7 – 266 mutations/Mb). A total of 14 cases (9.1%) were classified as TMB low (≤5 mutations/Mb), 64 of 154 (41.6%) were TMB intermediate (>5 and ≤20 mutations/Mb), 40 of 154 (26.0%) were TMB high (>20 and ≤50 mutations/Mb) and 36 of 154 (23.4%) were classified as TMB very high (>50 mutations/Mb). *NRAS* mutant melanoma demonstrated significantly decreased overall survival on multivariable analysis (HR for death 2.95, 95% CI 1.13 – 7.69, p = 0.027, log rank test) compared with other TCGA molecular subgroups. Other factors correlated with decreased overall survival included age and ECOG score. Of the 116 patients in our cohort with available treatment data, 36 received combination dual ICI with anti-CTLA4 and anti-PD1 inhibition as first line therapy. Elevated TMB was associated with significantly longer progression-free survival following dual agent ICI (HR 0.26, 95% CI 0.07 – 0.90, p =0.033, log rank test).

**Conclusions and Relevance:** *NRAS* mutation in CMs correlated with significantly worse overall survival. Elevated TMB was associated with increased progression-free survival for patients treated with combination dual ICI, supporting the potential utility of TMB as a predictive biomarker for ICI response in melanoma.

## Introduction

Cutaneous melanoma (CM) is associated with Ras mutations that activate downstream mitogen activated protein kinase (MAPK) signaling.^1–4^ CM can accordingly be molecularly classified based on MAPK pathway molecular driver which includes *BRAF* mutations, *NRAS* mutations and *NF1* deficiency, accounting for ∼80% of tumors.^2,4^ *BRAF, NRAS* and *NF1* are generally thought to be mutually exclusive oncogenic drivers although co-occurring driver mutations can be seen, particularly with *NF1* loss.^4^ *BRAF* and *NRAS* mutant melanomas histopathologically correlate with low cumulative sun damage (low-CSD) melanomas while *NF1* mutant melanomas are classified as high-CSD.^5^ The remaining 20% of melanomas are deemed “triple wild type” (TWT) and are clinically comprised of less common melanoma subtypes such as acral, mucosal and uveal melanomas occurring in sun-shielded locations (classified as non-CSD melanomas) and often driven by alternate pathways of melanomagenesis.

While driver group is associated with distinct demographic and pathologic features^4,6–8^, the relationship between molecular driver, clinical outcomes, and ICI response remains unclear. Prior large scale genomic studies including the TCGA landscape study of 333 primary and metastatic melanomas showed no significant correlations between genomic classification and clinical outcome.^4^ However, this was published prior to widespread ICI use, only one year after approval of anti-PD1 agents in metastatic melanoma.^9–12^

Therapeutic advances in melanoma have resulted in FDA approval of immune checkpoint inhibitors and several generations of BRAF targeted therapies for patients with *BRAF* V600E mutant tumors. Accordingly, *BRAF* mutant melanomas exhibit improved overall survival compared with other cutaneous melanomas.^13^ *BRAF* mutation was independently associated with improved recurrence free survival (RFS) in patients with stage 3 melanoma who were treated with adjuvant pembrolizumab.^14^ In addition, patients with *BRAF* mutation were shown to have superior outcomes when treated with first line ICI compared with BRAF and MEK inhibition and this has now become standard practice for a majority of patients.^15^ Improved survival outcomes in *BRAF* mutant melanomas thus may be related to disease biology in addition to the availability of multiple therapeutic avenues.

Pre-clinical work and retrospective genomic studies have implicated *NRAS* mutation as an overall poor prognostic factor with *NRAS* mutations associated with increased Breslow thickeness.^16,17^ However, data correlating clinical outcomes and response to ICI for *NRAS* mutant melanoma are mixed.^16,18–20^ The phase III IMspire170 trial compared cobimetinib + atezolizumab to pembrolizumab in patients with *BRAF* wild type melanoma and found no differences in PFS or ORR based on *NRAS*, *NF1* or TWT molecular driver in either of the two study arms.^21^ In a phase II trial evaluating ipilimumab vs ipilimumab/nivolumab, *NRAS* mutations were noted to be enriched in patients who experienced clinical benefit,^18^ suggesting molecular driver disease biology may affect ICI response in this patient population. With regard to other biomarkers of ICI response, the only robustly associated tumor-specific molecular feature is tumor mutational burden (TMB).^22^ High TMB correlates with improved ICI response and overall survival across a range of cancer types, including melanoma.^23^ Given almost all cutaneous melanomas have an intermediate or high TMB, it remains unclear whether there exists a TMB threshold above which the predictive significance of this biomarker becomes less robust.

Taken together, there remains a critical need to determine the precise relationship between molecular-genetic tumor alterations and clinical outcomes with contemporaneous systemic therapy regimens.^24^ Here, we retrospectively identified a real world cohort of 254 patients with CM undergoing targeted DNA sequencing of recurrently mutated cancer to evaluate whether any tumor specific genomic features, particularly TCGA driver and tumor mutational burden, correlated with clinical outcomes in the real-world setting. We additionally assessed for alternate mutations in TWT sun-exposed cutaneous melanomas to better understand this under-studied and poorly understood cutaneous melanoma phenotype. We found *NRAS* mutation was associated with poorer overall survival (OS), and TMB was predictive of dual ICI response, suggesting tumor genetics may help serve as an informative biomarker predictive of ICI response and clinical outcome for cutaneous melanoma.

## Methods

### Cohort Selection/Identification

A total of 330 patients with melanoma who underwent in house targeted next generation DNA sequencing were retrospectively identified using the UCSF tumor registry, from which medical record, baseline demographic, pathologic, and clinical outcome data were extracted.

Uveal, acral, mucosal, primary CNS/meningeal and pediatric melanomas were excluded. Melanomas of unknown primary (n =68) were assumed to be predominantly cutaneous and included, leading to a total of 254 cases included for subsequent analysis.

### Outcomes

The primary outcome assessed in this retrospective cohort was overall survival from the time of initial melanoma diagnosis. Secondary outcomes included progression-free survival evaluated in the context of systemic therapy for metastatic disease and recurrence-free survival for patients with localized disease treated with surgical intervention +/- adjuvant therapy. For the purposes of this study, progression free survival was defined as the time from initiation of first line systemic therapy to disease progression or death from any cause. Recurrence free survival was defined as time from date of primary resection for localized disease to the time of recurrent melanoma or death from any cause. Patients who developed recurrent disease and then subsequently received systemic therapy were included in both recurrence free survival and progression free survival analysis. Detailed information regarding systemic therapy regimen and timing for metastatic disease was available for 116 patients (Table 1). A majority of these patients received immunotherapy as first line systemic therapy (n = 108, 93.1%) with 8 patients receiving first line BRAF/MEK inhibition. Outcomes were assessed separately for patients who received first line anti-PD1 (n=65) and for patients who received first line combination dual ICI (anti-CTLA4 + anti-PD1, n = 36) as well as for all patients who received any form of immunotherapy in the first-line metastatic setting (including rare cases of anti-CLTA4 monotherapy, n =7).

**Table 1.**
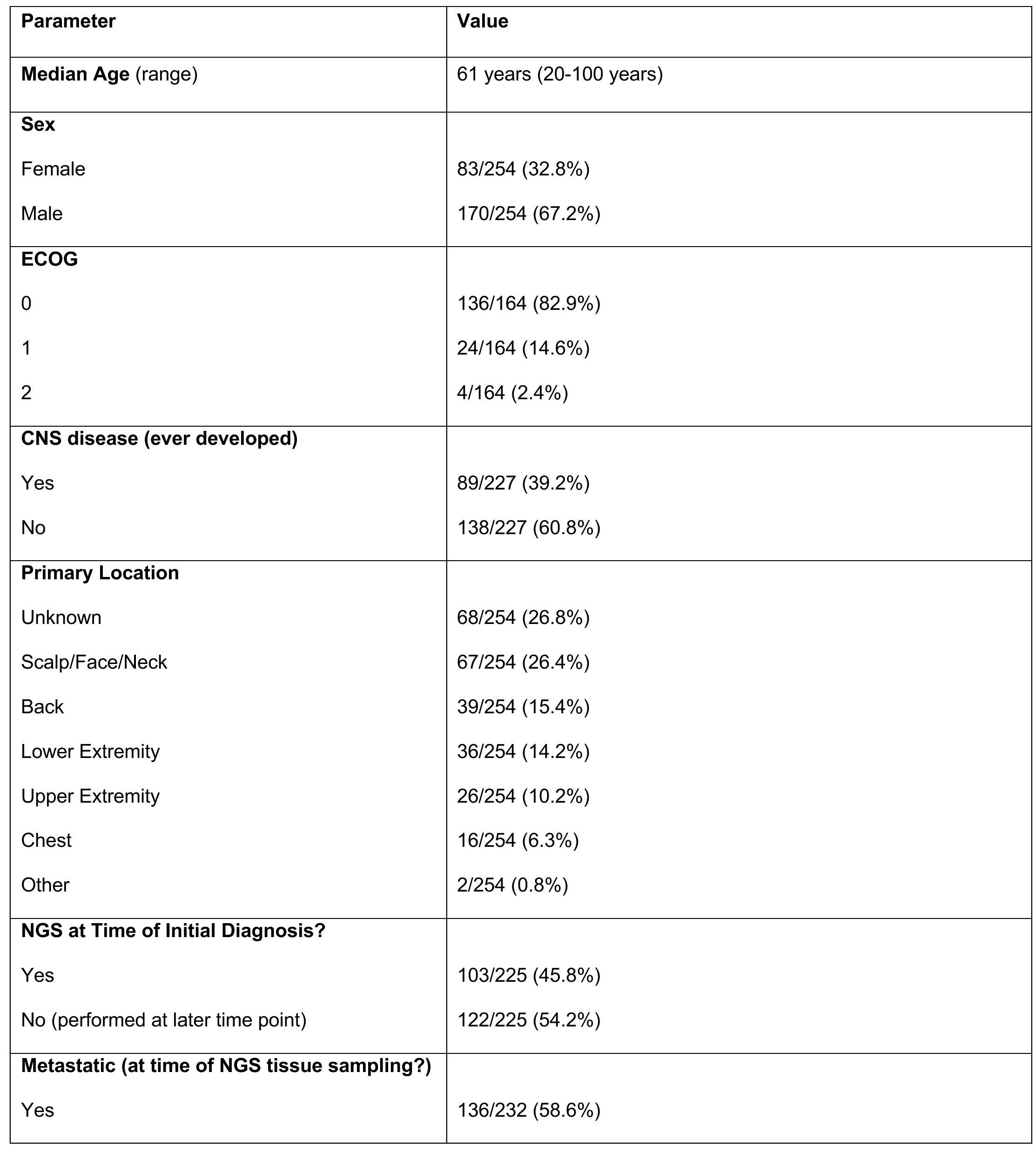
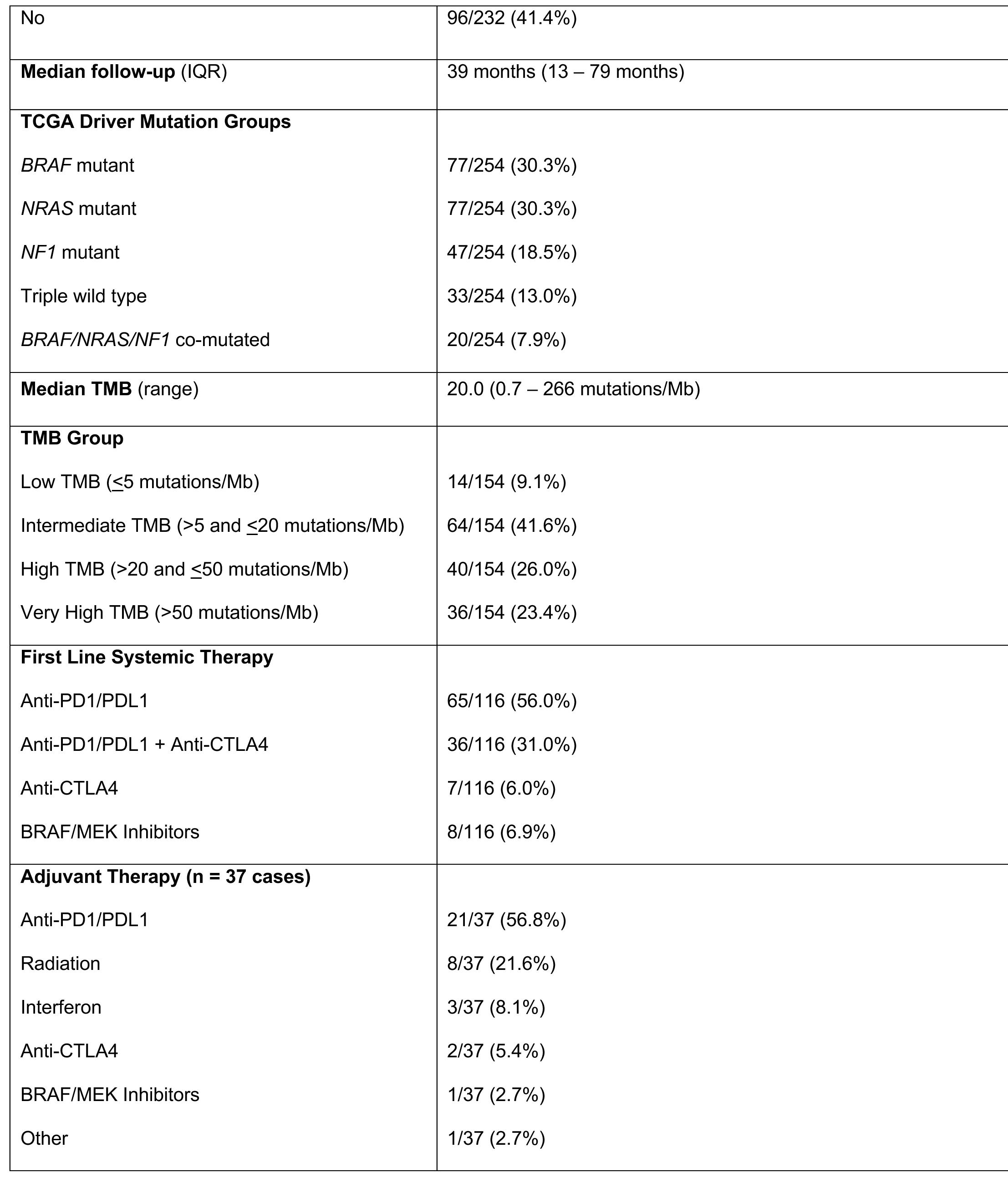
Baseline clinical characteristics of patients with metastatic melanoma undergoing target DNA sequencing (n=254 total patients).

### Statistical Analysis

Evaluation of categorical variables such as sex, ECOG status, stage, TMB group, TCGA driver gene was performed using either the chi-squared or Fisher exact test. Fisher’s exact test was used to assess proportions within two categorical variables. A chi-squared test was used to assess proportions within three or more categorical variables. An unpaired t-test was used for comparison of continuous variables such as age, TMB and MSI (%). All hypothesis tests were 2-sided and considered significant at α <0.05. Univariable Cox Proportional Hazards (CPH) was performed using STATA to evaluate overall survival, progression free survival and recurrence free survival and the following variables: age, sex, TMB, MSI (%), ECOG, presence of CNS disease, stage, TCGA mutation driver group (assessed independently for *BRAF, NRAS, NF1*, TWT and co-mutation group), any *NF1* mutation and TERT mutation status. Any variables found to have p <0.1 in univariable CPH analysis were included in a multivariable CPH model for overall survival (OS), progression-free survival (PFS) and recurrence-free survival (RFS). Kaplan Meier curves were then generated for OS, PFS and RFS using PRISM.

### Co-Mutation Analysis

In an effort to uncover other genes that were commonly altered together in this patient cohort, a co-mutation analysis was also performed. Each gene was first examined for the frequency at which it was altered across all patients, and the resulting gene list was filtered to only include the top 5% of genes that were most commonly altered. The ensuing threshold of ≥40 alterations resulted in 30 candidate genes for co-mutation analysis. Patients were then examined for the presence of alterations in these 30 genes, and the associations between genes were subsequently examined via Pearson correlation analysis. P-values were adjusted for multiple comparisons using the Bonferroni correction method, and significance was defined as an adjusted p < 0.05. All co-mutation analysis was conducted in R (R Foundation for Statistical Computing), version 4.2.2.

## Results

### Molecular melanoma groups display characteristic phenotypic and demographic features

Baseline demographic features for the 254 patients with cutaneous melanoma who underwent targeted DNA sequencing are summarized in Table 1. Targeted sequencing was performed at the time of initial melanoma diagnosis in 103/225 (45.8%) of cases and performed at a later time point in clinical care for melanoma in 122/225 cases (54.2%). A majority of cases (136/232, 58.6%) were metastatic at the time of tissue sampling for next generation sequencing. The median age for the overall cohort was 61 (range 20-100). Patients were predominantly male (n = 170/253, 67.2%), consistent with previously reported demographics for cutaneous melanoma in the United States.^25^ The median follow-up time for all patients was 39 months (IQR 13-79 months). A total of 186 cases were confirmed cutaneous melanoma (73.2%) while a total of 68 cases (26.8%) were melanoma of unknown primary (MUP). MUPs were included in our analysis based on histopathologic review by board certified dermatopathologists and prior data that suggests a majority of MUP exhibit mutational profiles consistent with a sun-exposed cutaneous origin (Table 1).^26,27^

Cases were classified based on previously reported TCGA driver genes including *BRAF* mutation (n=77, 30.3%), *NRAS* mutation (n=77, 30.3%), *NF1* mutation (n=47, 18.5%) and TWT (n= 33, 13.0%) (Figure 1a).^4^ A small cohort of cases demonstrated mutations in multiple TCGA driver genes (n=20, 7.9%), and within this subset, a majority harbored mutations in *NF1* as well as either *BRAF* or *NRAS* (n= 18 of 20, 90%) (Figure 1a). In addition to TCGA driver genes (*BRAF, NRAS, NF1*), we evaluated recurrent molecular alterations observed in at least 5% of the entire cohort, the majority of which are established oncogenes or tumor suppressors (Figure 1b).^28–41^ Recurrently altered genes included *TERT* (n=216), *CDKN2A/B* (n=116) *TP53* (n=93), *ARID2* (n=32), *PTEN* (n=25), *RASA2* (n=22), *PTEN* (n=25), *NFKBIE* (n=20), *KMT2B* (n=16), *PTPRD* (n=15), *PTPRT* (n = 15), *MAP2K1* (n=14), *RAC1* (n=14), *GRIN2A* (n=13), *PTPRB* (n=13 or 5%), *PREX2* (n=13 or 5%), and *SETD2* (n=13 or 5%).

**Figure 1.**
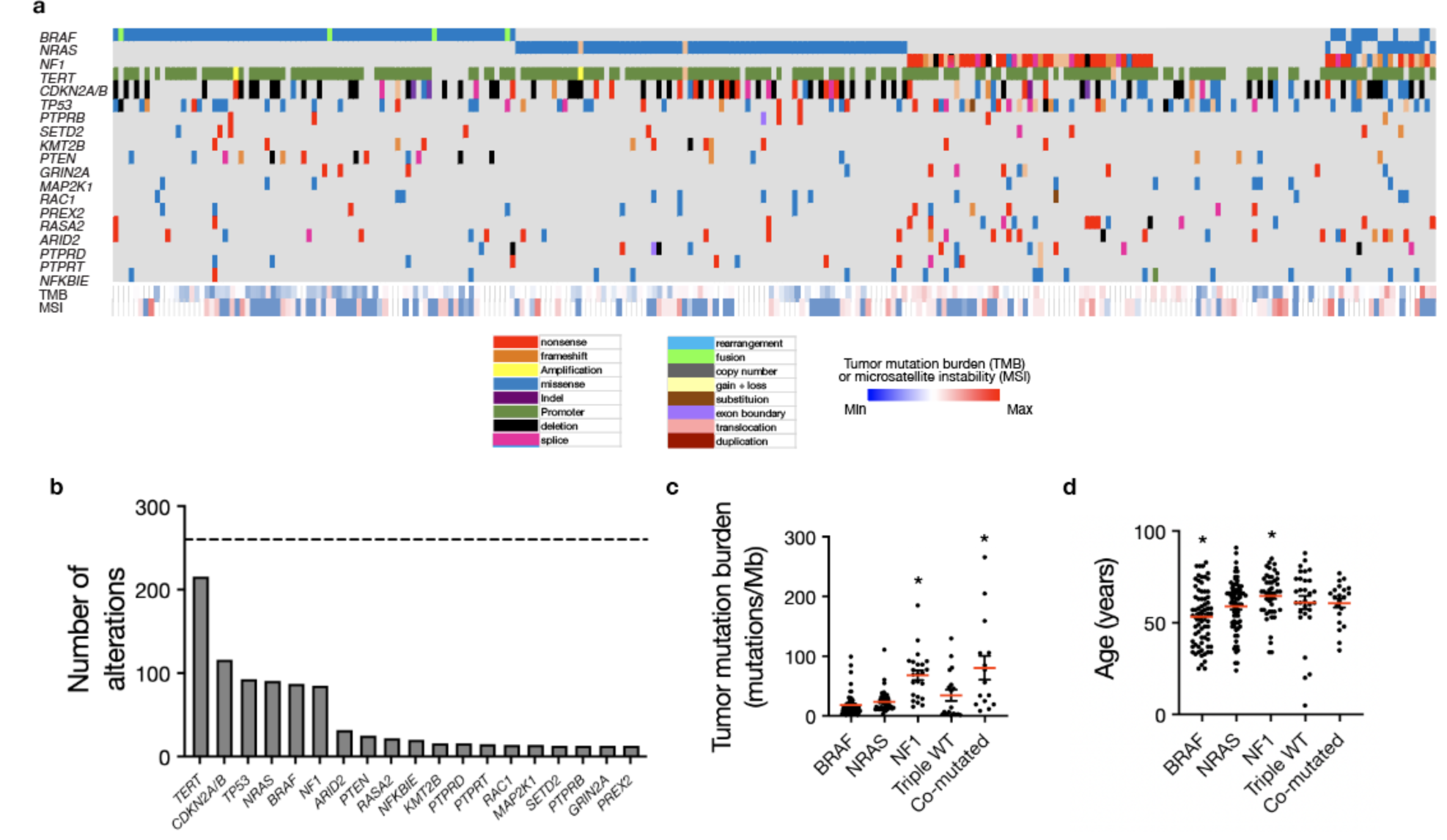
Mutational analysis of metastatic melanoma from a real-world cohort recapitulates TCGA molecular drivers. a. Targeted DNA sequencing analysis using a CLIA certified clinical genomics assay of 254 patients undergoing clinical care at a tertiary cancer center reveals five molecular groups: *BRAF* mutant (n=77), *NRAS* mutant (n=77), *NF1* mutant (n=47), triple wildtype (n=33), and co-mutated tumors with multiple alterations in *BRAF*, *NRAS*, and/or *NF1* (n=20) consistent with TCGA molecular groups. b. A total of 19 genes were recurrently altered in at least 5% of the cutaneous melanoma cohort including *TERT* (n=216), *CDKN2A/B* (n=116), *TP53* (n = 93), *NRAS* (n = 91), *BRAF* (n=87), *NF1* (n=85), *ARID2* (n = 32), *PTEN* (n = 25), *RASA2* (n=22), *NFKBIE* (n = 20), *KMT2B* (n =16), *PTPRD* (n = 15), *PTPRT* (n=15), *RAC1* (n=14), *MAP2K1* (n=14), *SETD2* (n=13), *PTPRB* (n=13), *GRIN2A* (n=13), and *PREX2* (n=13). c. Tumor mutation burden is significantly increased in *NF1* mutant and co- mutated tumors compared to *BRAF, NRAS,* or TWT tumors (ANOVA, p<0.001). d. Patients with *BRAF* mutant tumors are significantly younger while *NF1* mutant patients are significantly older than other molecular groups (ANOVA, p=0.001).

*BRAF* mutant melanoma tended to occur in younger patients with a median age of 54 (p=0.0001, one-way ANOVA), which may account in part for previous reports of improved outcomes in *BRAF* mutant melanoma, while *NF1* mutant tumors were seen in older patients with a median age of 67 (p=0.005, one-way ANOVA) compared with other driver genes (Figure 1d). Tumor mutational burden was significantly higher in *NF1* mutant tumors (median 70.2 mutations/Mb) and co-mutated tumors (median 66.5 mutations/Mb) (p<0.001, one-way ANOVA) compared with other TCGA driver genes (Figure 1c). *BRAF* mutant tumors had the lowest median TMB (12.4 mutations/Mb,) followed by *NRAS* mutant tumors (median 17.5 mutations/Mb) and TWT melanomas (28.2 mutations/Mb) (Figure 1c). Patients with *BRAF* mutant tumors were associated with increased rates of non-metastatic disease at diagnosis (p<0.0001, chi-square test) while patients with TWT tumors were associated with poorer performance status (ECOG 1+) (n=7/20, 35%) compared with other TCGA driver genes (n=21/144, 14.6%; p = 0.028, Fisher’s exact test) (Figure S1c-d). With regards to anatomic location, *NF1* mutant tumors (n = 22/47, 46.8%), TWT melanomas (n=14/33, 42.4%) and co-mutant tumors (n=7/20, 35%) occurred more frequently on the head/neck compared with *BRAF* or *NRAS* mutant tumors (n = 24/154, 15.6%; p<0.001, chi-squared test) (Figure S1a). There were otherwise no statistically significant differences in baseline features between TCGA driver genes (Table 1; Figure S1b,e,f).

BRAF, NRAS, *and* NF1 *mutant melanomas exhibit distinct alteration and co-mutation patterns. BRAF* mutations can be classified into three groups: Class I, II or III.^42^ Class I and Class II mutations lead to Ras independent Raf activation, with Class I mutations involving V600 to create monomers with high kinase activity and Class II mutations creating activated Raf dimers with intermediate kinase activity. In contrast, class III mutations comprise weaker oncogenic variants leading to enhanced Ras binding with impaired kinase activity that drives CRAF activation for enhanced downstream MAPK pathway signaling.^43^ In our cohort, *BRAF* mutations were predominantly class I (Figure 2a), particularly for the *BRAF* driver group, in which 68 of 77 cases (88.3%) had class I *BRAF* mutations. Only one class III *BRAF* mutation was noted in this group (*BRAF* G466E) with several class II mutations.^42^ Multiple fusions and gene rearrangements were also detected involving *AGK* and *BRAF* (n=3) and *ATF7* and *BRAF* (n=1). In contrast, only one of the 11 *BRAF* mutations identified in the *BRAF/NRAS/NF1* co-mutation group was a class I *BRAF* mutation. When occurring in the co-mutation group, a majority (8 of 11 or 73%) of *BRAF* mutations were class III, consistent with prior work suggesting these weaker oncogenic *BRAF* variants require additional hits to drive melanomagenesis.^42^

**Figure 2.**
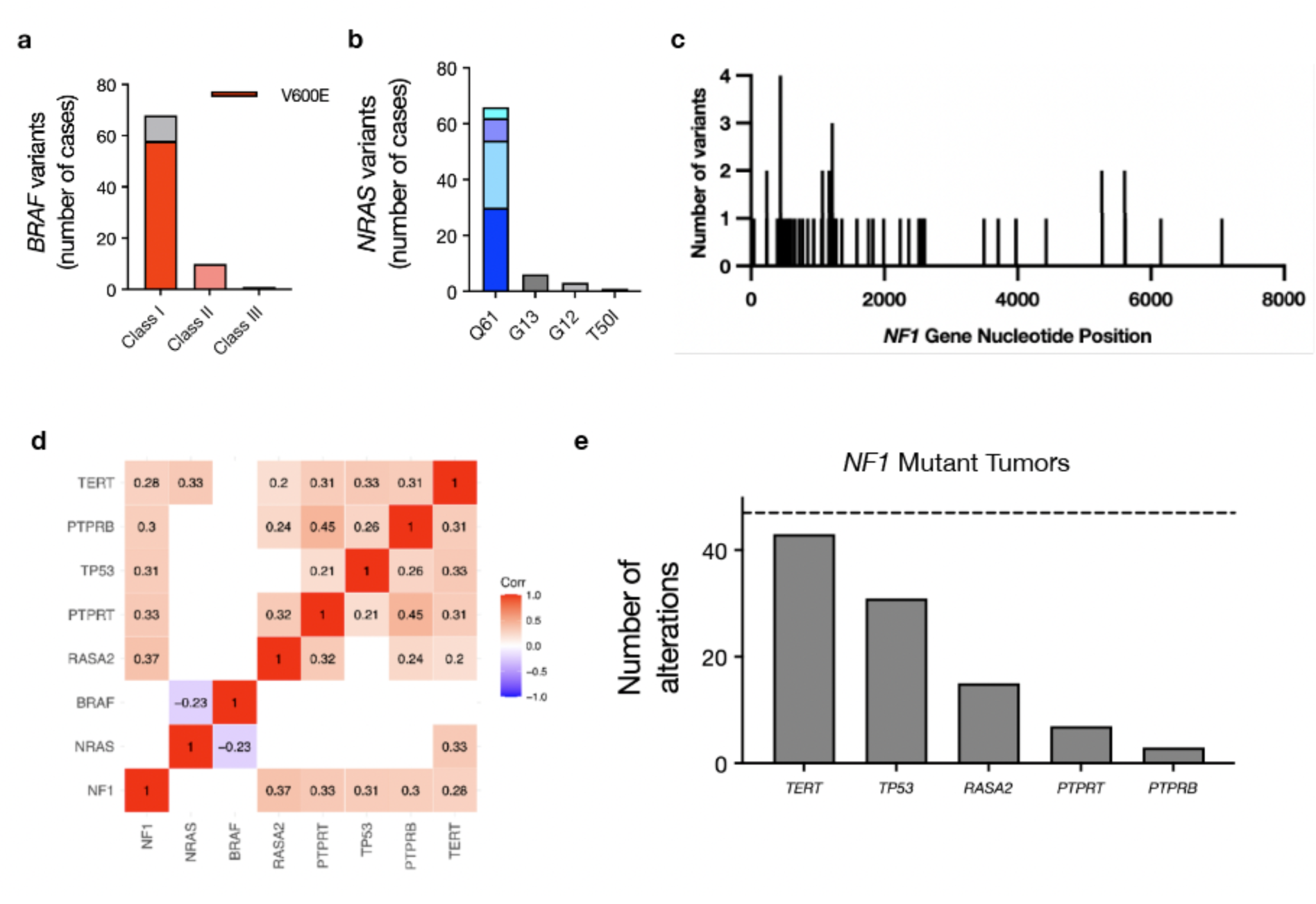
Driver mutation specific analysis of BRAF, NRAS and NF1 mutant tumors reveals specific variant and co-mutation patterns. a. *BRAF* mutant tumors were primarily composed of class I *BRAF* mutations (n = 68 or 86%). The majority were V600E (n=58) with 8 V600K mutations and 2 V600R mutations. A total of 3 *BRAF* fusions were detected as well as 2 gene re-arrangements, most of which involved *AGK* and *BRAF*. Other non-class I loci in this group included K601, G469, G466, F247 and E586. There were no associated differences in outcomes with class I mutations compared to other *BRAF* classes or to non-*BRAF* mutant tumors. Only 1 of 10 *BRAF* mutant cases with co-occurring *NRAS* or *NF1* mutations (10%) had a class I *BRAF* mutation. S467L mutations were particularly common in this cohort (n=4 or 40%). Other loci included N581, G469, L597 and N580. b. A majority of *NRAS* mutant tumors had mutations at Q61 (N =66 or 87%) including Q61R (n = 30), Q61K (n=24), Q61L (n=8) and Q61H (n=4). Other mutations identified included G13R (n=5), G12A (n=3), G13C (n=1), and T50I (n=1). Two amplifications were noted as well as one case with copy number neutral LOH at Q61K. Of the 15 *NRAS* mutations identified in the co-mutation group, 10 were Q61K/L/R (67%), 2 were G12A/D, 2 were G13D and one was T50I. c. A wide spectrum of variants, enriched on the 5’ end of the gene, was identified in *NF1* mutant tumors with 12 of 47 cases in the *NF1* driver group (26%) and 8 of 18 *NF1* mutant cases (44%) in the *BRAF/NRAS/NF1* group having more than one mutation in *NF1*. The presence of multiple *NF1* mutations did not have a significant impact on overall survival when compared with tumors with only one *NF1* alteration. A majority of the *BRAF/NRAS/NF1* co-mutant tumor group had mutations in *NF1* (n=18 or 90%). There were no significant survival outcome differences between this group and the *NF1* driver mutation group. d. Co-mutation correlation analysis (threshold Pearson > 0.3 at p<0.05) found that *BRAF* mutations did not significantly correlate with any other alterations, *NRAS* mutations correlated with *TERT* mutations and *NF1* mutations were significantly correlated with several genes including *RASA2*, *PTPRT*, *TP53*, *PTPRB* and *TERT*. Notably, *BRAF* and *NRAS* mutation were anti-correlated consistent with their mutual exclusivity as drivers. e. Co-mutation breakdown of *NF1* mutant tumors and significantly co-occurring alterations.

*NRAS* mutations were predominantly Q61 variants regardless of whether tumors were in the *NRAS* driver (n=67/77 or 86%) or co-mutated group (n=9/14 or 64.3%) (Figure 2b). A total of 67 putative loss of function variants in NF1 were identified, with the majority clustered at the 5’ end of the gene, suggesting upstream loss of function mutations are enriched in *NF1* (Figure 2c). There were no differences in specific *NF1* variants when co-mutated with other driver genes compared to tumors with *NF1* mutation alone.

Correlation analysis of all identified genes found *NF1* mutations to correlate with several genomic alterations including most frequently *RASA2* loss (correlation coefficient 0.37, adjusted p < 0.001) but also frequently with *PTPRT* mutations (correlation coefficient 0.33, adjusted p < 0.001), *TP53* mutations (correlation coefficient 0.31, adjusted p < 0.001), *PTPRB* mutations (correlation coefficient 0.30, adjusted p < 0.001) and 39 other less commonly co-mutated genes (Figure 2d-e; Figure S2). *NRAS* correlated most with *TERT* (correlation coefficient 0.33, p < 0.001), *BRAF* was not significantly co-altered with any genes examined, and *BRAF* and *NRAS* demonstrated significant mutual exclusivity (Figure 2d, Figure S2).

### Triple wild type tumors contain frequent alterations in other components of MAPK pathway signaling

A majority of TWT tumors demonstrated a pathogenic variant in the Ras pathway (n=28/33 or 84.8%) (Figure 3a-b). These included *MAP2K1* (n =7), *SPRED1* (n=4), *KIT* (n=3), *FGF4/19* (n=3), *MAP3K8* (n=2), *RASA2* (n=2) and *ERBB2* (n=2), *MET* (n=2), *NF2* (n=2) and *GAB2* (n=2). (Figure 3b, 3d). In addition, triple wildtype tumors often harbored non-Ras pathway alterations found in other TCGA molecular groups including *TERT* (n=21)*,TP53* (n=17), *CDKN2A/B* (n=12), *NFKBIE (n=3)*, *YAP1 (n=3)*, *FAT1 (n=3)*, *NOTCH1 (n=3)*, *NOTCH2 (n=3)*, *PTEN (n=3)*, *MITF (n=3)* and *ATM (n=3)* (Figure 3c, d). A total of 5 of 33 TWT cases demonstrated no known or suspected Ras pathway alterations and demonstrated presumed driver mutations in *EIF1AX*, *GNAQ*, *ATRX* and *ASXL1* (Figure 3a).

**Figure 3.**
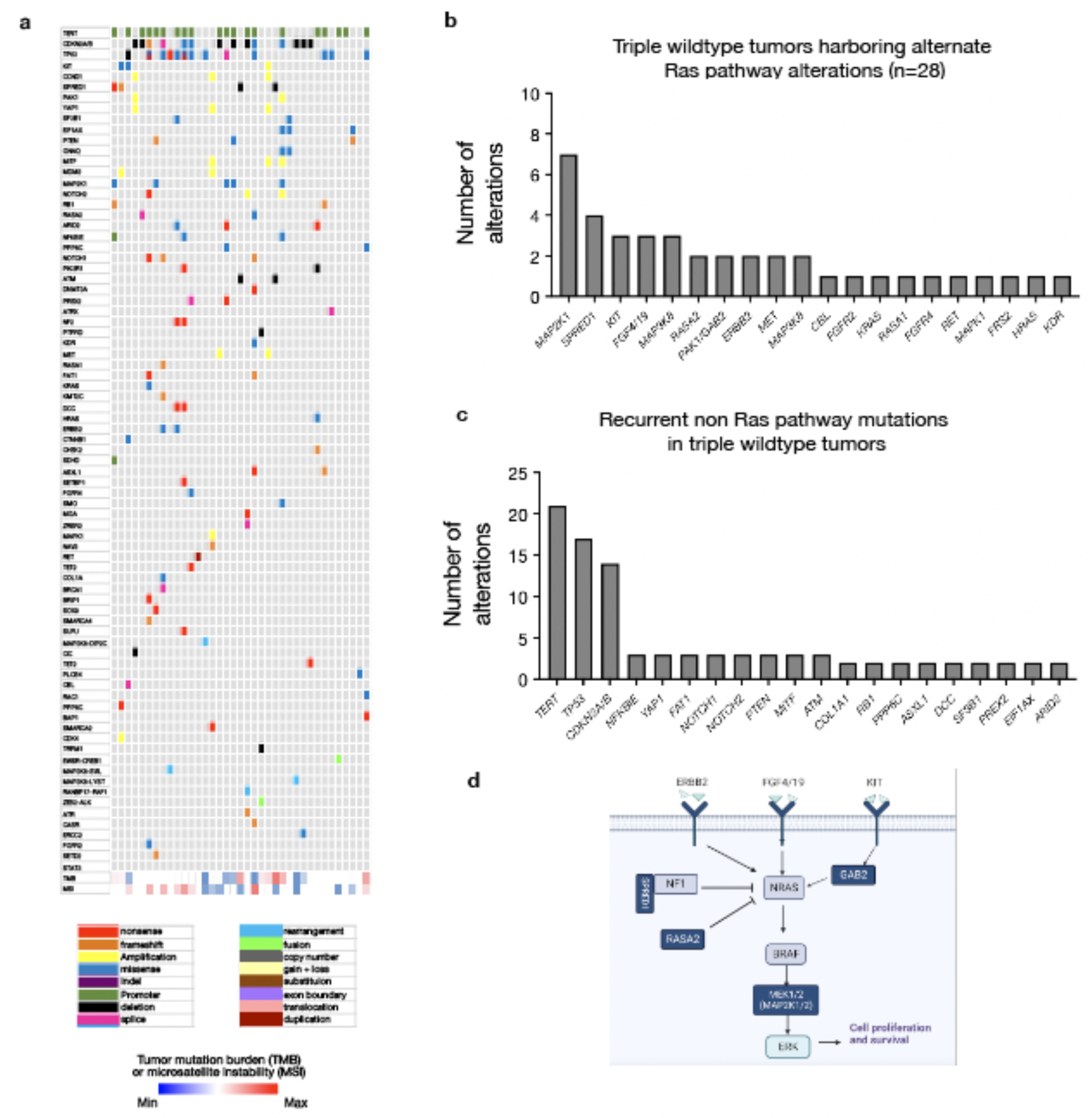
Triple wild type tumors harbor alternate Ras pathway alterations in the majority of tumors lacking a classical TCGA driver mutation. a. In triple wildtype tumors, recurrently altered genes included *TERT* (n=21), *TP53* (n=18), *CDKN2A/B* (n=14), *MAP2K1* (n=7), and *SPRED1* (n=4). b. In 28 of 33 cases (84.8%), pathogenic variants in known Ras pathway components were observed as putative driver mutations. c. Recurrent alterations in non-Ras pathway genes in the triple wildtype group are predominantly *TERT* and *TP53* mutation similar to other molecular groups, with additional mutations comprising potential drivers in tumors without a classic Ras pathway mutation. d. Schematic depicting mutations occurring in triple wildtype tumors (dark blue) within the Ras signaling pathway.

### NRAS mutation is prognostic, and tumor mutational burden (TMB) is predictive of response to dual checkpoint blockade

We next assessed clinical outcomes for the entire 254 patient cohort. With regard to overall survival (OS), univariable CPH analysis revealed age, ECOG status, CNS disease, stage, *BRAF* mutation and *NRAS* mutation were significantly correlated with OS (p<0.1; Table S2). On multivariable analysis, *NRAS* mutant tumors remained correlated with worse OS (HR 2.95, 95% CI 1.13 – 7.69, p=0.027; log-rank test) (Figure 4a,Table S2). Older age (HR 1.06, 95% CI 1.02 – 1.11, p = 0.003; log-rank test) and higher ECOG score (HR 4.84, 95% CI 2.12 – 11.05, p <0.001; log-rank test) also correlated with decreased overall survival (Figure 4b, Table S2). Multivariable analysis for recurrence free survival found age to be the only variable correlated with RFS (HR 1.03, 95% CI 1.02 – 1.05, p <0.001, log rank test) (Table S4). Of the 116 patients treated with systemic therapy for metastatic or locally advanced/unresectable disease, the majority were treated with first line ICI: 65 of 116 (56.0%) patients received anti-PD1 monotherapy, 36 of 116 (31.0%) patients received dual checkpoint inhibition with combination anti-PD1 and anti-CTLA4 and 7 of 116 patients (6.0%) received anti- CTLA4 monotherapy (Table 1). A small proportion were treated with BRAF/MEK inhibitors in the first line setting (n=8/116 or 6.9%). Multivariable analysis found no significant associations with progression free survival on any first line immune checkpoint inhibition (Table S3). When further stratified based on specific treatment type, TMB was the only variable significantly associated with response to dual ICI; no variables associated with response to single agent anti- PD1 (Figure 4c, Table S5-6). Low TMB (≤5 mutations/Mb) correlated with poorer PFS in patients receiving dual agent ICI (HR 20.9, 95% CI 1.77 – 245.9; p = 0.016; log-rank test) (Figure 4d, Table S6).

**Figure 4.**
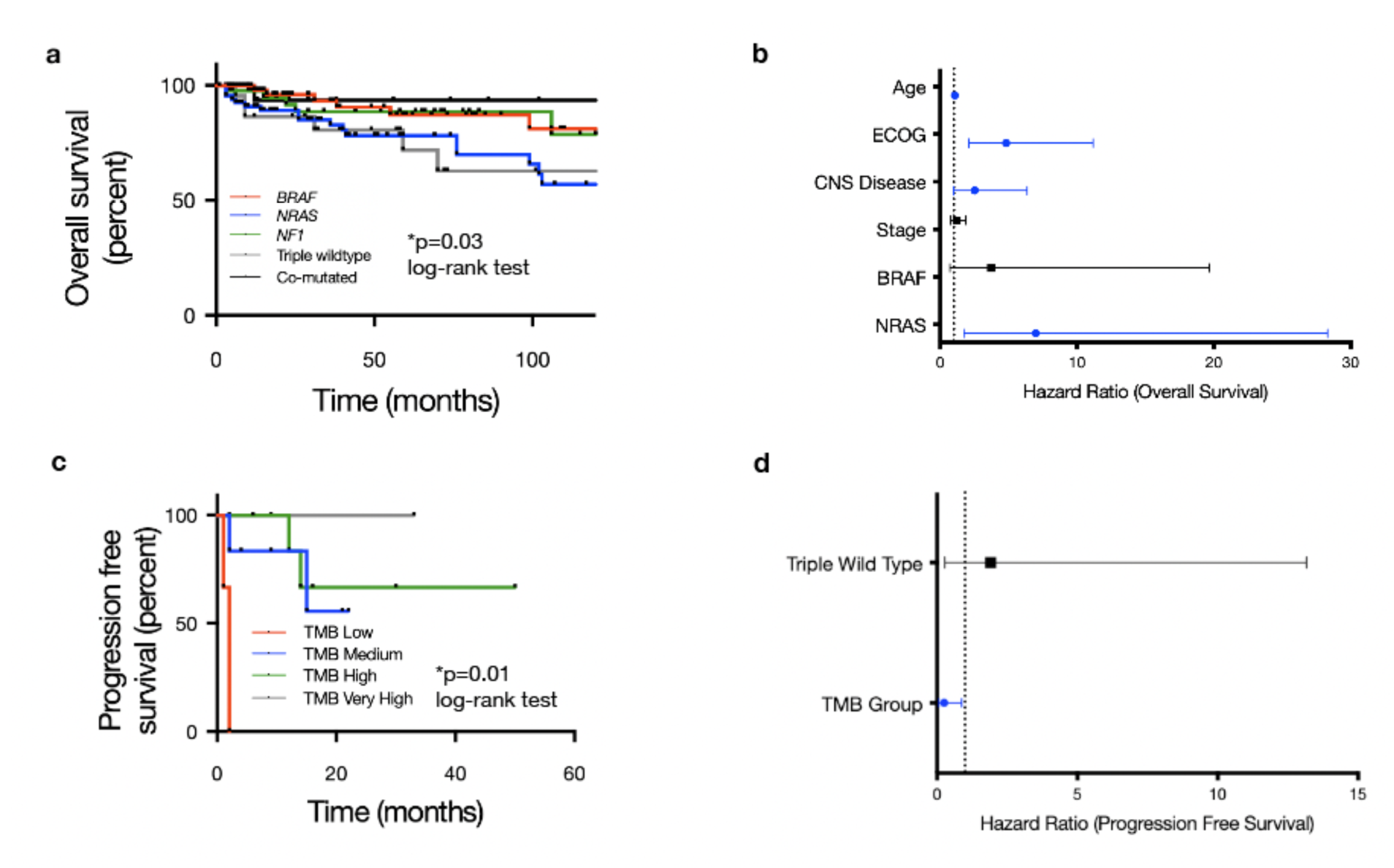
Molecular subgroup is prognostic for overall survival and tumor mutation burden (TMB) is predictive of dual checkpoint blockade response in cutaneous melanoma. a. Overall survival stratified by 5 pre-specified molecular subgroups (*BRAF* mutant, *NRAS* mutant, *NF1* mutant, triple wildtype and co-mutated *BRAF/NRAS/NF1*) shows *NRAS* mutation is associated with worse outcomes. b. Forest plot of multivariable hazard ratios for OS including covariates with p ≤0.1 in univariable survival analysis (ECOG status, presence of CNS disease, stage, BRAF, NRAF). In addition to *NRAS* mutation, age, ECOG status and the presence of CNS disease were independent factors significantly associated with overall survival. c. Kaplan Meier curve depicting progression free survival stratified by tumor mutation burden (TMB) for the 36 of 116 patients (31.0%) who received the combination of anti- PD1/PDL1 + anti-CTLA4 as first line systemic therapy for metastatic disease. d. TMB was the only covariate found to significantly impact PFS on combination ICI in multivariable CPH analysis.

## Discussion

We present clinical and molecular data from 254 patients with melanoma who underwent targeted DNA sequencing and were treated in the ICI era. Our analysis confirmed CM classification based on TCGA genetic drivers^4^ and previously reported associated demographic and phenotypic features including age of onset (*BRAF* mutations in younger patients and *NF1* mutations found in older patients) and significantly higher median TMB in *NF1* mutant tumors ^4,8^ (Figure 1c-d). *BRAF* mutations in the entire 254 patient cohort were predominantly class I (n = 68 of 88 or 77.3%) (Figure 2a). While cases with *BRAF* TCGA driver mutations alone were consistent with previously reported data suggesting a predominance of class I mutations in *BRAF* mutant melanoma, only one of eleven *BRAF* co-mutated cases (co-occurring with either *NRAS* or *NF1*) exhibited a class I *BRAF* mutation. The majority of co-mutated cases had weaker class III *BRAF* mutations, which have been associated with improved prognosis in melanoma compared with class I *BRAF* mutations.^44^ We found *NF1* to be the only TCGA driver in our cohort that significantly correlated with multiple other genes, most frequently *RASA2* as previously reported.^45^ *NF1* co-mutations with either *BRAF* or *NRAS* as well as frequently with other melanoma associated oncogenes suggests that this is overall a weaker oncogenic driver that may cooperate with additional MAPK pathway alterations to drive melanomagenesis. *NRAS* mutant tumors correlated with poorer overall survival compared with other TCGA driver gene groups. The presence of an *NRAS* mutation has historically been correlated with poorer overall survival in patients with advanced melanoma but its prognostic impact in the modern treatment era is less clear.^46,47^ Our analysis of patients treated in the immunotherapy era (from 2013- 2023) found *NRAS* mutation to be an independent factor significantly associated with poorer overall survival, underscoring the need for further drug development in this particular patient population.

The majority of TWT tumors carried at least one putative driver mutation involving the MAPK pathway, including *MAP2K1, SPRED1, KIT, FGF4/19, RASA2* and *ERBB2* (Figure 3). Triple wild type tumors also demonstrated a median TMB of 28.2, suggesting that the correlation between low TMB and triple wild type status may be most relevant to sun-shielded or non-cutaneous melanomas (Figure 1c). Only 5 of 33 TWT melanomas in our cohort lacked a putative driver mutation involving the MAPK pathway. Our analysis suggests the majority of TWT cutaneous melanomas harbor a Ras/MAPK pathway mutation with some confirmed cutaneous melanomas demonstrating mutations more commonly identified in uveal melanoma including *EIF1AX* and *GNAQ*.

Beyond TCGA drivers, several studies have focused on identifying additional prognostic or predictive tumor-specific genomic aberrations in cutaneous melanoma, particularly in terms of ICI response. Mutations in *KMT2*, *PTPRT*, *MAP2K1* and *SETD2* have been associated with improved responses to immune checkpoint inhibitors while *PTEN* loss of function is thought in some studies to be associated with innate resistance to immune checkpoint blockade.^48–52^ We did not find any specific genomic alterations other than the presence of an *NRAS* mutation to correlate with clinical outcomes in our 254 patient cohort. TMB was the only predictive biomarker of response to immune checkpoint inhibition identified in our analysis. Notably, TMB was not associated with PFS on single agent anti-PD1 but was predictive for patients treated with dual agent immune checkpoint inhibition in our patient population. Interestingly, despite a significantly higher TMB than other driver groups, *NF1* mutant and co-mutant tumors did not demonstrate improved PFS when treated with first line immunotherapy. Given higher TMB has been correlated with response to ICI, both in our study and in several others, and TMB in *NF1* mutant melanomas is generally very high, further investigation is warranted as to why *NF1* mutation alone did not correlate with improved ICI response.

## Conclusions

Our real-world retrospective analysis of 254 patients with melanoma who underwent in house DNA sequencing at our institution generated several novel and notable findings. *NRAS* mutant tumors were associated with worse overall survival outcomes. We also demonstrate that triple wildtype cutaneous melanomas that lack mutations in *BRAF*, *NRAS* or *NF1* have a similar TMB to the overall median and in a majority of cases demonstrate putative drivers in the MAPK pathway, suggesting that disease biology in these cases is likely similar to those melanomas with known Ras drivers. Finally, TCGA driver group was not associated with progression free survival on any first line immunotherapy despite prognostic significance in terms of overall survival. Tumor mutational burden was associated with response to combination immune checkpoint inhibition with anti-CTLA4 and anti-PD1 but did not have a statistically significant association with progression free survival on anti-PD1 monotherapy. Taken together, these data support prognostic differences based on driver mutation and predictive utility for TMB in cutaneous melanoma. Additional investigation into “triple wild type” cutaneous melanomas and their reliance on Ras/MAPK signaling is an area for further study, and poor outcomes in *NRAS* mutant tumors highlights an urgent, unmet clinical need for therapeutic development.

Our study has several limitations, most notably that it is a retrospective single institution experience, with all the caveats such an approach entails. In addition, detailed follow-up and treatment data were available only for a subset of the 254 patients that underwent molecular profiling. Further prospective studies that assess the prognostic significance of driver mutations and predictive and prognostic significance of tumor mutational burden are certainly warranted.

---

**Figure S1.**
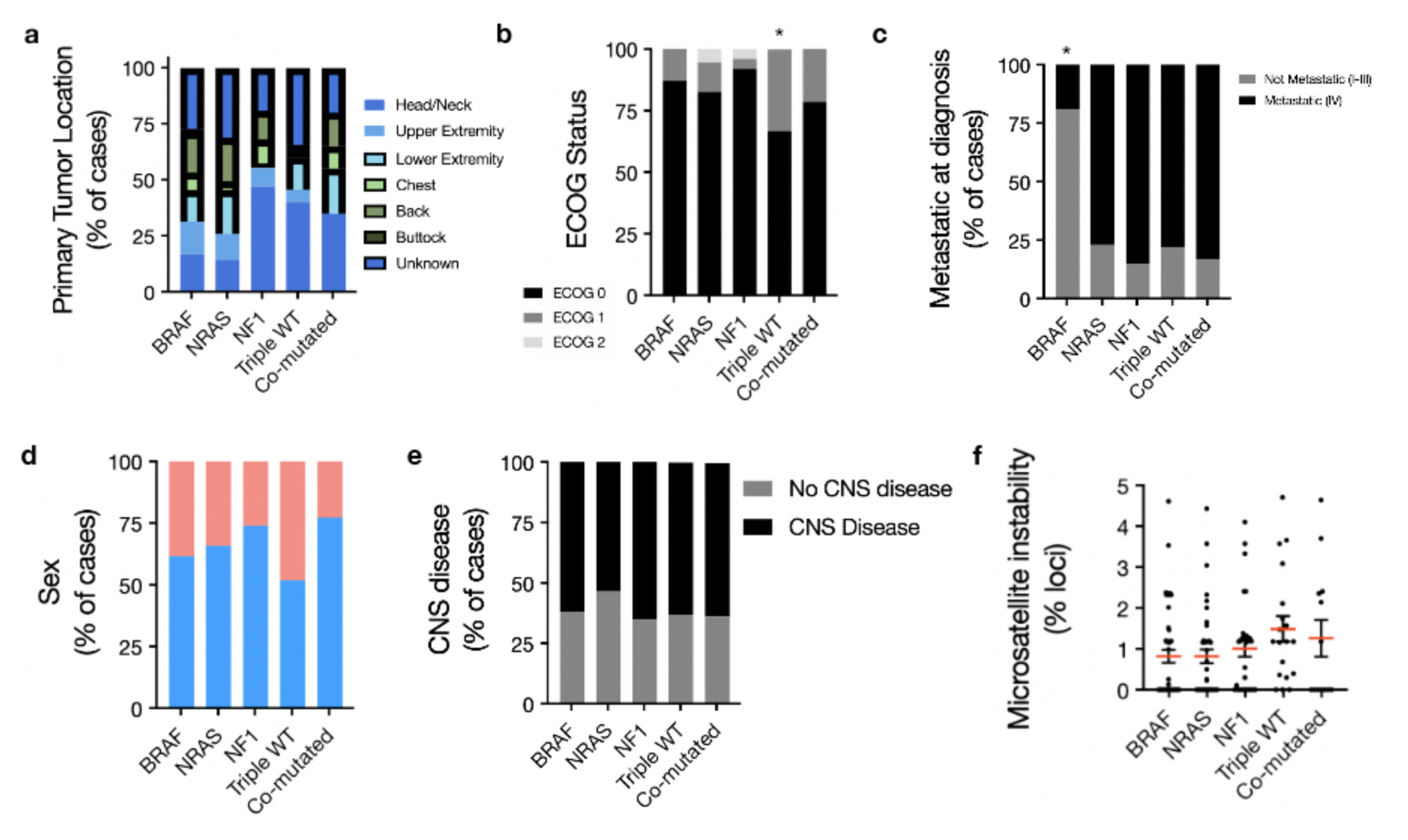
Baseline clinical correlations based on molecular group. a. Location of primary tumor based on molecular driver. *NF1* mutant, co-mutated and triple wild type tumors were more likely to occur on the head/neck. b. There were no statistically significant differences in sex breakdown between molecular subgroups. c. *BRAF* mutation was significantly associated with non-metastatic presentation at diagnosis (chi-square test p < 0.0001) . d. TWT tumors were significantly more likely to present as ECOG 1+ (n=7/21, 33%) compared with non TWT tumors (n=21/146, 15%).; p = 0.033. e. There were no statistically significant differences in rates of development of CNS disease between TCGA driver groups. f. There were no statistically significant differences in microsatellite instability (MSI) between molecular driver groups.

**Figure S2.**
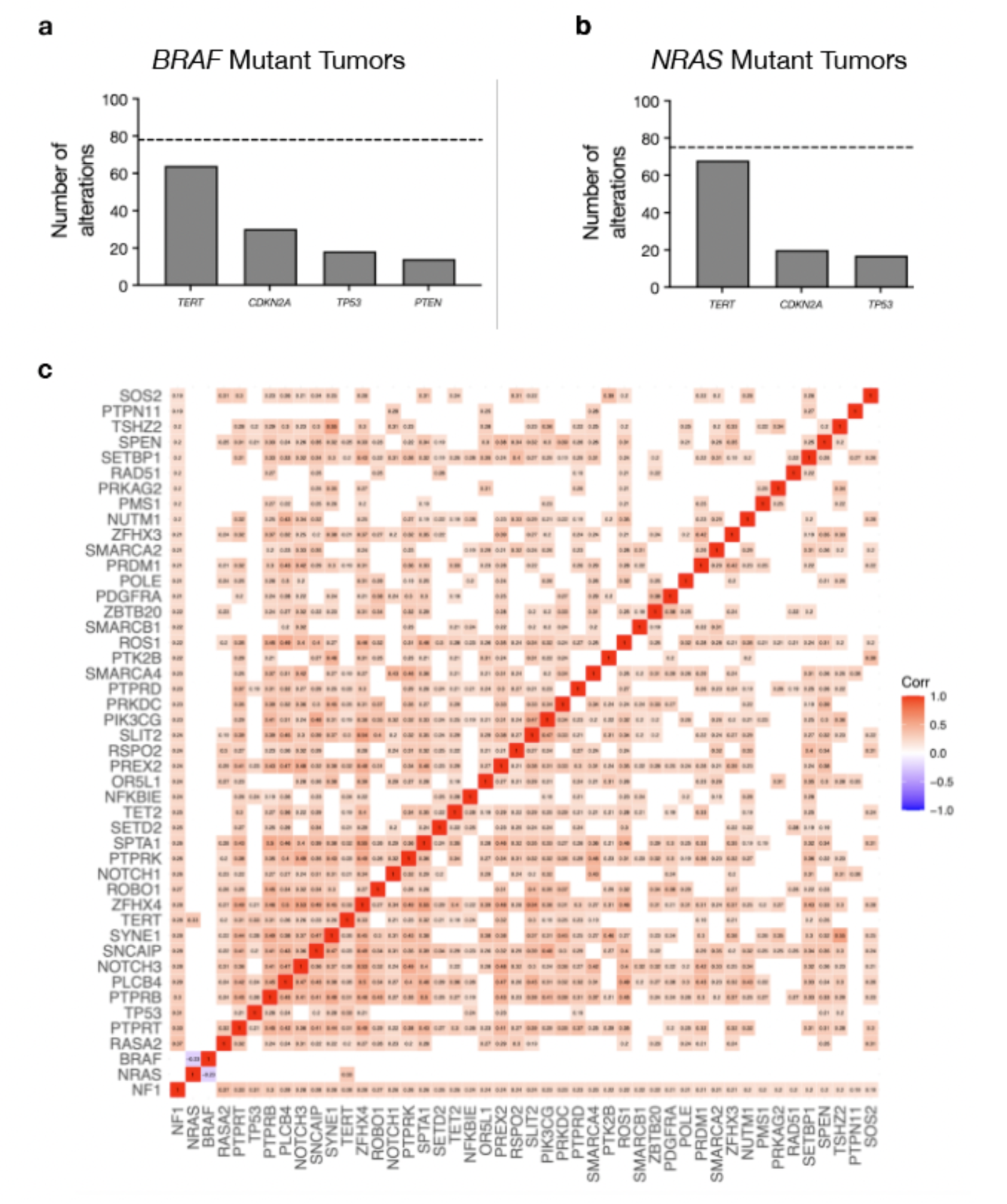
Analysis of co-mutation patterns. a. Most frequently co-altered genes in *BRAF* mutant tumors. b. Most frequently co-altered genes in *NRAS* mutant tumors. c. Correlation matrix summarizing statistically significant gene correlations across the entire cohort without imposing a strict Pearson correlation coefficient threshold.

**Figure S3.**
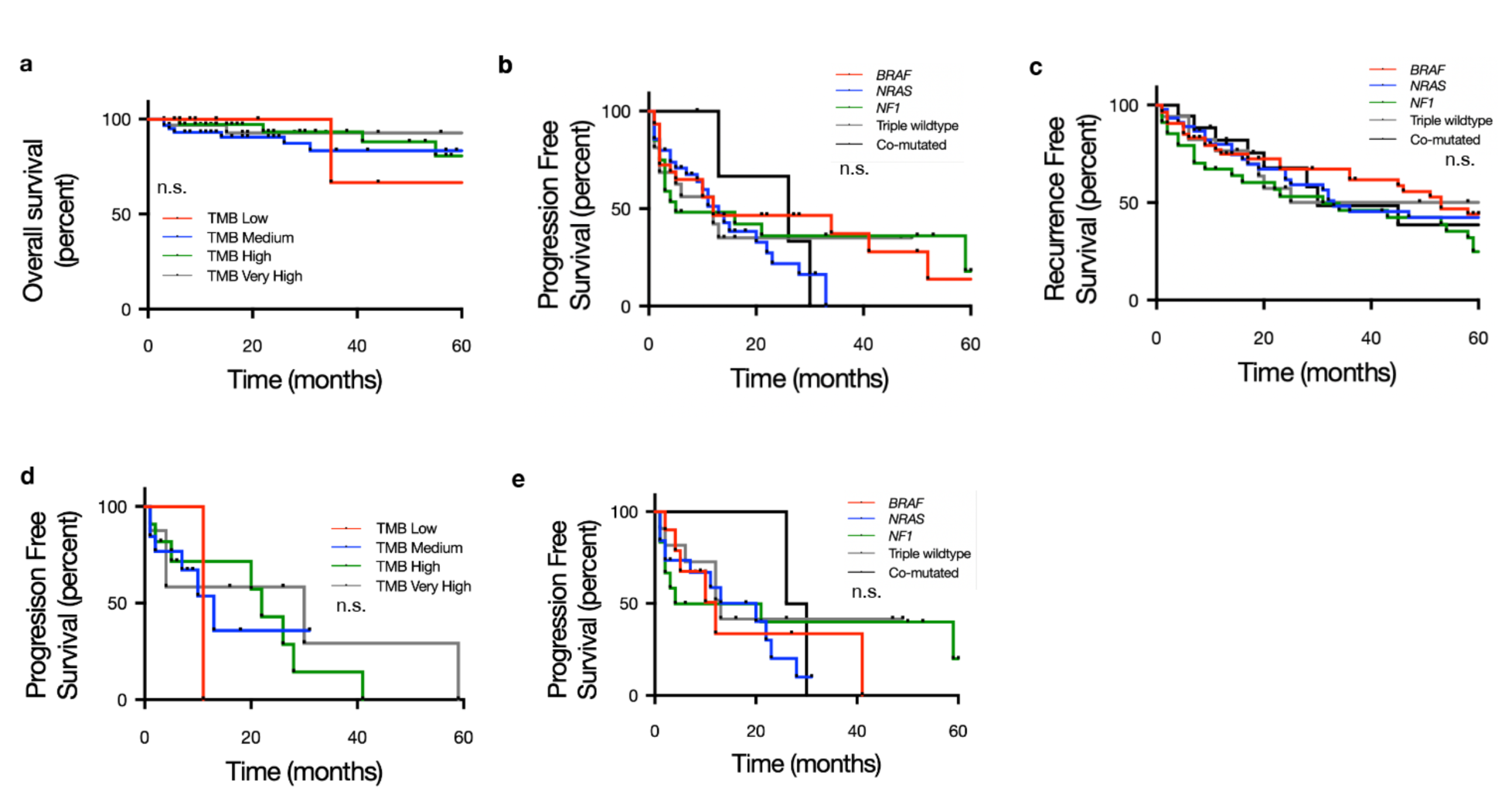
Clinical outcomes based on molecular signatures and treatment. a. Kaplan Meier (KM) curve depicting overall survival stratified by TMB group which showed no statistically significant differences in overall survival outcomes for all 254 patients included. b. KM curve depicting progression free survival on any first line systemic therapy based on TCGA molecular driver which showed no statistically significant differences between groups. c. KM curve depicting recurrence free survival following resection based on TCGA molecular driver. There were no statistically significant differences in RFS between driver groups. d. KM curve depicting progression free survival on anti-PD1 monotherapy stratified by TMB group. There were no statistically significant differences in PFS on anti-PD1 between these groups. e. KM curve depicting PFS on anti-PD1 monotherapy stratified by TCGA molecular driver. There were no statistically significant differences in PFS on anti-PD1 monotherapy between TCGA driver groups.

**Supplementary** Table 1. Compiled variant list from the UCSF500 assay, a CLIA certified, capture based targeted DNA sequencing assay of 529 cancer-associated genes.

**Supplementary** Table 2. Univariable and multivariable analysis for OS.

**Supplementary** Table 3. Univariable and multivariable analysis for PFS on any first line immunotherapy.

**Supplementary** Table 4. Univariable and multivariable analysis for RFS

**Supplementary** Table 5. Univariable and multivariable analysis for PFS on single agent immune checkpoint inhibition.

**Supplementary** Table 6. Univariable and multivariable analysis for PFS on dual agent immune checkpoint inhibition.

